# Human cortex organizes dynamic co-fluctuations along sensation-association axis

**DOI:** 10.1101/2025.07.14.660681

**Authors:** De-Zhi Jin, Changsong Zhou, Xi-Nian Zuo, Joshua Faskowitz, Ting Xu, Ye He

## Abstract

The human brain dynamically organizes its activity through coordinated fluctuations, whose spatiotemporal interactions form the foundation of functional networks. While large-scale co-fluctuations are well-studied, the principles governing their amplitude-dependent transitions—particularly across high, intermediate, and low-amplitude regimes—remain unknown. We introduce a co-fluctuation score to quantify how instantaneous functional interactions reorganize with global amplitude dynamics. Using resting-state fMRI data, we identified amplitude-dependent co-fluctuation transitions between functional systems hierarchically aligned with the sensorimotor-association (SA) axis: sensorimotor networks dominated high-amplitude co-fluctuations, associative systems prevailed during intermediate amplitudes, and limbic system preferentially engaged in low-amplitude states. This hierarchy underwent developmental refinement from childhood to adulthood and adaptively reconfigured under external stimuli. Replicated across four independent samples including 7T fMRI, these findings establish the SA axis as infrastructure for amplitude-dependent transitions of co-fluctuation states. Our framework bridges transient coordination and stable functional architecture, demonstrating how brain networks balance external processing (high-amplitude states) with internal cognition/emotion (mid-to-low amplitude states) through amplitude-stratified interactions.

## Introduction

Spontaneous fluctuations in blood-oxygen-level-dependent (BOLD) signals measured by functional magnetic resonance imaging (fMRI) have been a central focus for researchers studying brain function. These fluctuations exhibit temporally coherent activity across different brain regions, a phenomenon termed functional connectivity (FC), which has become a widely used metric to study functional interactions across the brain ^1,2^. The spatial organization of whole-brain FC underpins functional systems or networks related to diverse processes, including visual, sensorimotor, attention functions and etc ^3^. These networks can support complex behaviors and cognitive processes ^4^, and are reproducible across individuals ^5,6^.

While static FC provides valuable insights, it fails to capture the dynamic nature of brain activity. Emerging evidence highlights that brain regions interacting in complex and time-varying patterns with brain activity fluctuates dynamically. These transient configurations of FC reveal rich dynamical structures in functional networks at fine timescales ^7–10^. Methods such as sliding-window correlations and coactivation patterns (CAPs) have been employed to characterize these dynamic interactions ^7,10–14^. Time-varying characteristics of brain connectivity have been observed under awake and anesthetized conditions ^15^, and are associated with cognitive performances ^16–19^ and clinical status ^20,21^. At the system level, brain networks exhibit hierarchical dynamics ^22^, with periodic pattern of segregation and integration emerging over time ^23,24^. However, methods such as sliding-window and CAPs are limited by their inability to resolve single-frame patterns and their reliance on thresholding to filter low-amplitude moments.

A recent method focusing on co-fluctuation timeseries has enabled the investigation of functional interaction fluctuations at a single-frame timescale ^25–27^. This method interprets FC as a summary of co-fluctuations in activities between brain regions over time ^28^, making it particularly suited for capturing transient changes in functional interactions ^29^.

Studies utilizing this approach have primarily examined high-amplitude co-fluctuation moments (events), which are thought to drive static FC ^26^, and are linked to individual brain fingerprints ^30^, cognition ^31^ and movie-watching states ^29,32^. However, this analysis as well as CAPs focus on high-amplitude events, considering it as “true” signals, while frequently dismissing low-amplitude signals as susceptible to noise. Low-amplitude co-fluctuation moments can also capture critical aspects of brain network information ^33–35^. This raises critical questions: what co-fluctuation patterns underpin and shape different co-fluctuation amplitudes? How do these time-varying co-fluctuation patterns support functional networks? Addressing these questions is essential for advancing our understanding of the intrinsic functional organization of the brain.

In this study, we decomposed FC into frame-wise co-fluctuation patterns to explore the time-varying functional organization of human cerebral cortex across different global co-fluctuation amplitudes. We propose a novel metric, the co-fluctuation score, defined as the ratio of regional to global co-fluctuation amplitude (Fig. 1). This metric quantifies relative co-fluctuations in individual regions compared to whole-brain co-fluctuation, allowing for unbiased comparisons across amplitudes. Our findings reveal amplitude-dependent co-fluctuation transitions between functional systems hierarchically aligned with the sensorimotor-association (SA) axis: sensorimotor networks dominated high-amplitude co-fluctuations, associative systems prevailed during intermediate amplitudes, and limbic system preferentially engaged in low-amplitude states. These patterns exhibited a gradual maturation trajectory from childhood to adulthood and were modulated during movie-watching states. We replicated the main findings in independent datasets including a 7T fMRI dataset. These results suggest that spontaneous brain activities interact within a structured, amplitude-dependent framework that hierarchically transit from sensorimotor to association dominance.

**Fig. 1.**
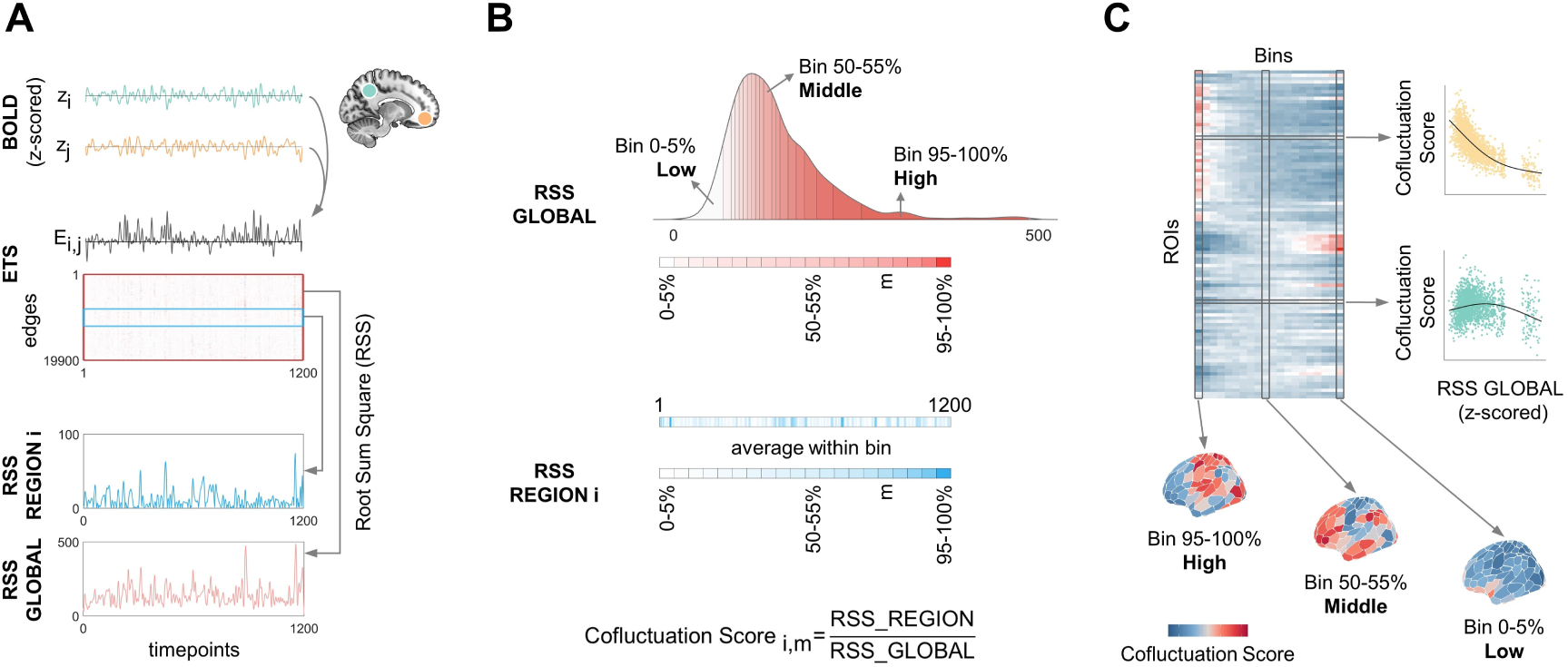
Schematic illustration for amplitude-stratified co-fluctuation analysis. (**A**) Computing edge-wise co-fluctuation amplitudes. For each pair of brain regions, their edge timeseries (ETS, E_i,j_ ) is derived from the element-wise product of z-scored BOLD timeseries (z_i_ and z_j_). At each timepoint, global or regional co-fluctuation amplitudes are computed as the root sum of squares (RSS) of ETS across all edges (RSS GLOBAL) or across edges connected to a given region (RSS REGION). (**B**) Amplitude-stratified co-fluctuation scoring. Timepoints are ranked by their global RSS values and partitioned into 20 bins (5 % of timepoints/bin). Within each bin, global and regional RSS values are averaged. The regional co-fluctuation score is defined as the ratio of regional RSS to global RSS from a specific bin. (**C**) Two analytical strategies of co-fluctuation scores. First, model amplitude-dependent trajectories via generalized additive models. Second, compare spatial patterns across global co-fluctuation amplitude bins.

## Results

FC can be unwrapped into a timeseries of co-fluctuations between a pair of brain regions at each timepoint (Fig. 1A). Global co-fluctuation amplitude (RSS GLOBAL) measures the instantaneous co-fluctuation level across the cerebral cortex. It is not directly related to the global signal (fig. S1). Co-fluctuation score quantifies the relative co-fluctuation level of a given region with respect to the whole-brain, allowing for unbiased comparisons across amplitudes (Fig. 1B). We calculated co-fluctuation scores for each of the 200 brain regions defined by Shaefer parcellation ^36^ for each subject from 100 unrelated subjects of the Human Connectome Project (HCP) dataset (Fig. 1C, see Materials and Methods for details).

### Hierarchies of amplitude-stratified co-fluctuation dynamics across cortical networks

To better understand how regional co-fluctuations dynamically change under different global co-fluctuation amplitudes, we delineated the trajectories of co-fluctuation scores changing with global amplitudes by employing Generalized Additive Models (GAMs) for each brain region (Fig. 2A).

**Fig. 2.**
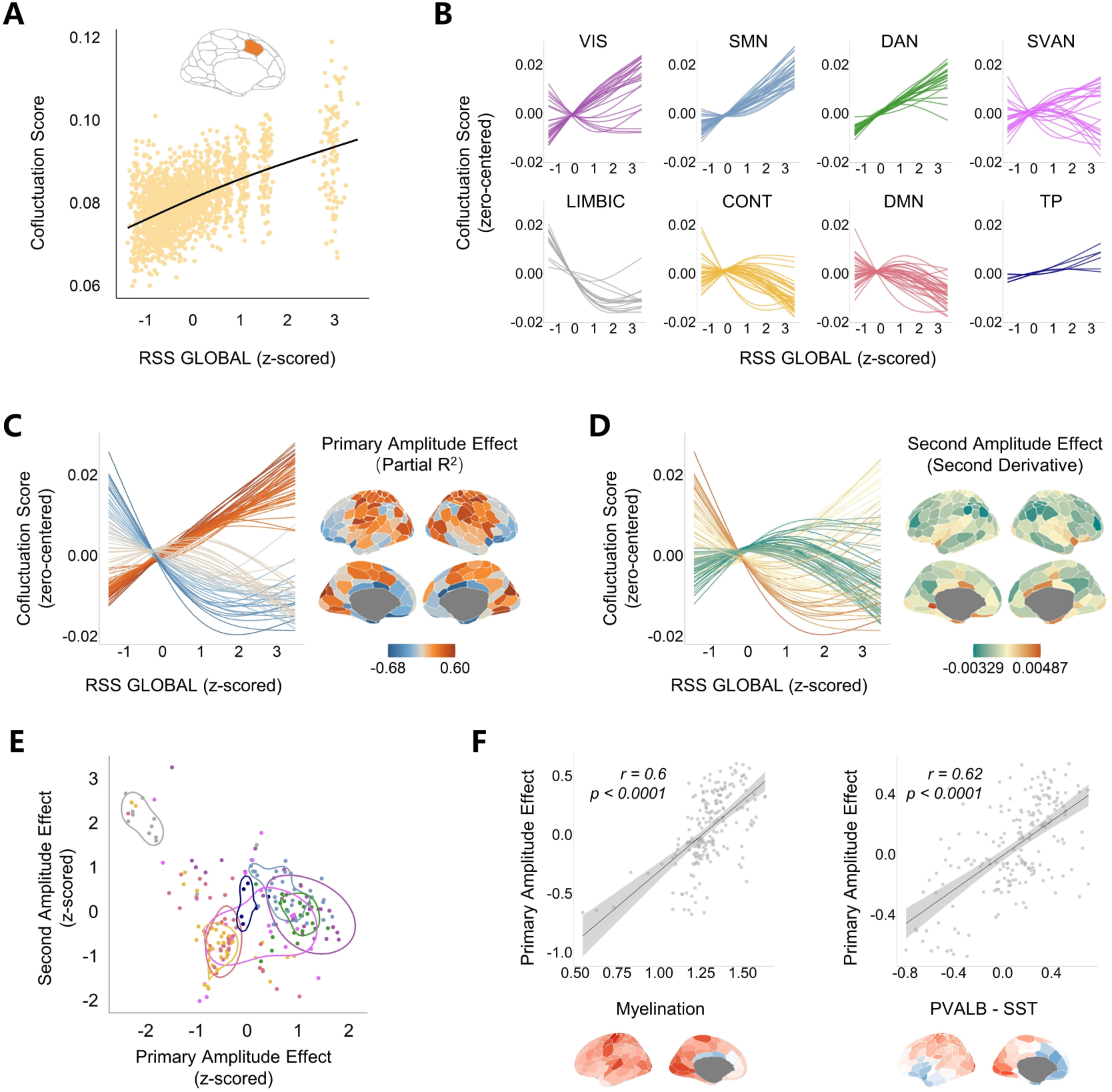
Amplitude-dependent trajectories of co-fluctuation scores. (**A**) Amplitude-dependent co-fluctuation score dynamics for a representative region of 100 subjects modeled via generalized additive models (GAMs), adjusted for sex and mean framewise displacement. (**B**) Co-fluctuation score trajectories were grouped for eight canonical functional networks (VIS-visual, SMN-somatomotor, DAN-dorsal attention, SVAN-salience/ventral attention, LIMBIC-limbic, CONT-control, DMN-default mode, TP-temporal parietal). (**C**) Primary amplitude effects. Trajectories of regional co-fluctuation scores are colored by partial R^2^, with each line representing a single brain region. (**D**) Second amplitude effects. Trajectories of regional co-fluctuation scores are colored by the averaged second-order derivatives. Only the top and bottom 50 regions for both the primary and second amplitude effects are shown to clearly depict their distinct trajectories. (**E**) Scatter plot of the primary and second amplitude effects colored by eight functional networks. The lines represent the contours of the highest density regions of each network generated by two-dimensional density estimation with ggdensity package in R. (**F**) Primary amplitude effects correlate with T1w/T2w myelination map (left) and the difference map of normalized parvalbumin and somatostatin (PVALB-SST) expression (right).

From low to high global amplitude, the changing patterns of co-fluctuation scores showed systematic variations across eight canonical functional networks (Fig. 2B). Regions within somatomotor (SMN), visual (VIS) and dorsal attention (DAN) networks exhibited near-linear increases in co-fluctuation scores with rising global amplitudes. The majority of regions in control (CONT) and default mode (DMN) networks followed inverted U-shape trajectories. Regions within limbic network (LIMBIC) showed a downward trend as global co-fluctuation amplitudes increased. In addition, regions within ventral attention network (SVAN) exhibited mixed profiles, including both upward and inverted U-shape patterns. These patterns delineate three amplitude-dependent coupling regimes: the primary sensorimotor systems driving global co-fluctuations through linear amplification; higher-order cognitive systems balancing co-fluctuation with peaks under the intermediate amplitudes; and limbic systems engaging more during low-amplitudes states and decoupling during high-amplitude states.

We quantified these effects through partial R^2^ (linear) and the averaged second derivatives (quadratic) of the trajectories, namely the primary and second amplitude effects (Fig. 2, C and D). The orthogonal distribution of the primary and second amplitude effects (Fig. 2E) revealed hierarchical gradients of amplitude-dependent co-fluctuation reorganization, extending from the limbic regions to sensorimotor (mean centile: x = 74.8%, y = 60.8%) or higher-order systems (mean centile: x = 32.1%, y = 32.1%). It further confirmed three types of amplitude-dependent co-fluctuation reorganization. The primary and second amplitude effects may reflect a spatial-temporal organizing principle of dynamic functional architecture.

We further explored the correlates of regional heterogeneity of the primary amplitude effects with neuroanatomical and transcriptomic profiles. Previous works suggest cortical myelination and the relative presence of parvalbumin (PVALB) and somatostatin (SST) reflect hierarchical functional differences across cortex ^37–40^. Their regional heterogeneity may be related to the amplitude effects. Consistent with this prediction, significant positive correlations of the primary amplitude effects with cortical myelination (r = 0.6, pspin < 0.0001; Fig. 2F), as well as with relative densities of PVALB-SST (r = 0.62, pspin < 0.0001; Fig. 2F) were observed.

### Amplitude-dependent spatial configuration of co-fluctuation scores along the SA axis

To further explore how the spatial organization of regional co-fluctuation scores changed with respect to different global co-fluctuation amplitudes, we computed group-averaged spatial maps of co-fluctuation scores for each amplitude bin (Fig. 3A, the full set of 20 brain maps is shown in fig. S2). Distinct spatial configurations emerged across global amplitudes. High-, intermediate-, and low-amplitude bins revealed spatial patterns of sensorimotor, associative and limbic systems respectively. The spatial similarities between these maps revealed two clusters with dissociation between high-amplitude bins and mid-to-low amplitude bins (Fig. 3B). Neurosynth meta-analysis further demonstrated functional segregation of these states. High-amplitude maps of co-fluctuation scores predominantly correlated with sensorimotor function, while intermediate-amplitude maps aligned with high-order cognitive domains like memory and language. Low-amplitude maps mainly linked to social and emotion functions (Fig. 3C).

**Fig. 3.**
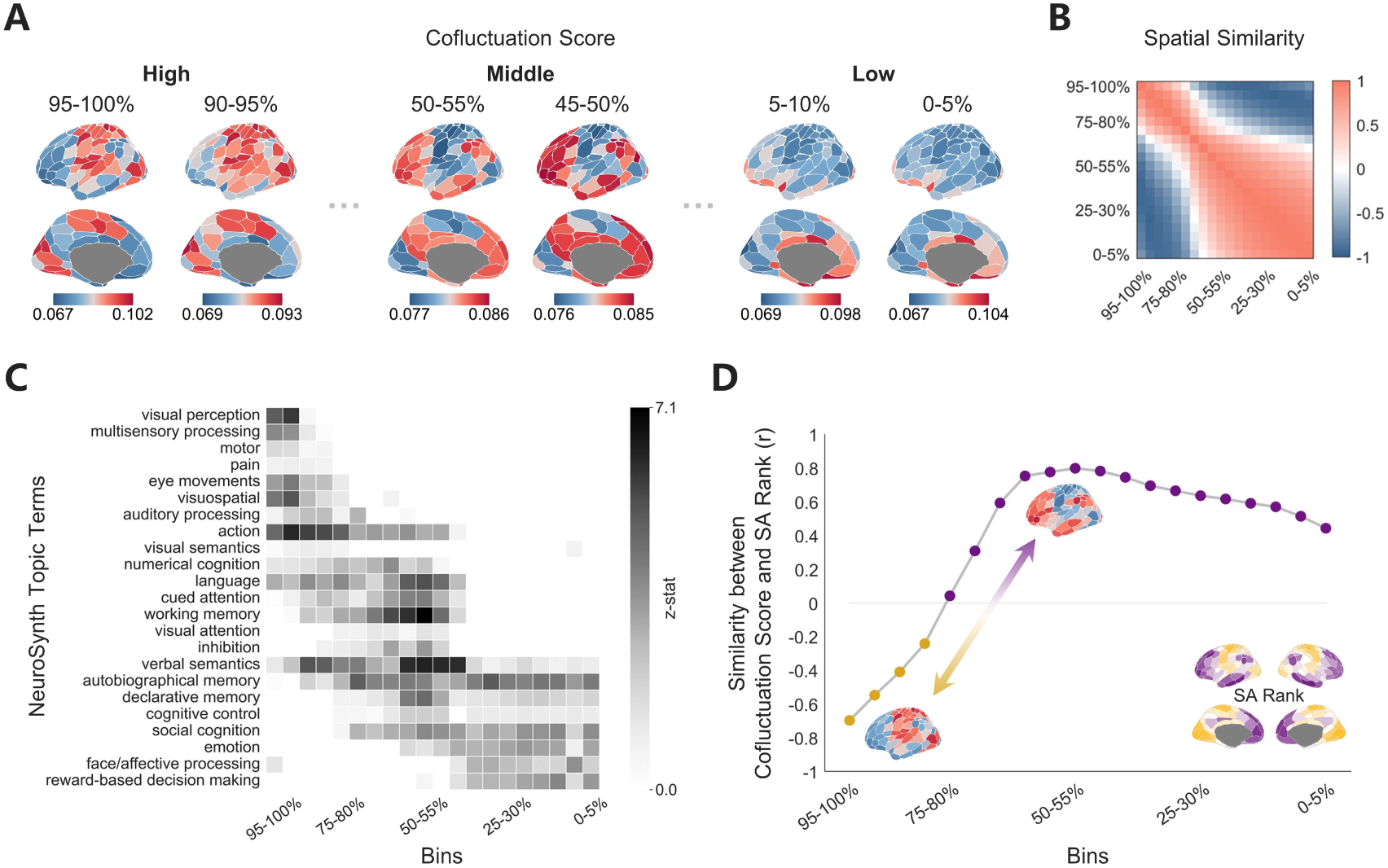
Amplitude-dependent spatial configuration of co-fluctuation scores. (**A**) Group-averaged maps of co-fluctuation scores for high, intermediate and low amplitude bins. Sensorimotor regions dominate high-amplitude states, whereas associative regions peak at intermediate amplitudes, and limbic system prevail during low-amplitude states. (**B**) The spatial similarities of co-fluctuation score maps across 20 bins (Pearson’s correlation coefficients), revealing two clusters. (**C**) Neurosynth meta-analysis of top 20% regions per bin decoded with 23 topic terms identifies amplitude-stratified functions. (**D**) Sensorimotor-association (SA) axis alignment. Spearman’s correlation between co-fluctuation score maps and SA-rank map transitions from high amplitudes to low amplitudes.

Next, we assessed the spatial alignment of these spatial maps with the sensorimotor-association (SA) axis, which represents a fundamental hierarchy of human cortex followed by multiple neurobiological properties ^41–44^. As illustrated in Fig. 3D, the spatial similarities between co-fluctuation score maps and the SA axis exhibited systematic amplitude dependence. Notably, the two identified clusters exhibited opposing patterns along the SA axis. The highest-amplitude map (top 5% bins) of co-fluctuation scores was strongly aligned with the sensorimotor-dominated SA axis (r = -0.69, pspin < 0.0001). This relationship weakened progressively with decreasing amplitude, reaching non-significance at bin 75-80%. Beyond this turning point, the relationship reversed, with co-fluctuation score maps increasingly aligning with the association-dominated SA axis and peaked at mid-amplitudes (at bin 50-55%, r = 0.79, pspin < 0.0001). These amplitude-dependent shifts suggest that dynamic co-fluctuation across amplitudes is constrained by the SA axis, with switching between three systems: sensorimotor regions driving global co-fluctuation at high amplitudes, association regions engaging at mid-amplitudes, and limbic regions prevailing during low-amplitude states. This dynamic alignment may underscore the hierarchical and amplitude-dependent organization of brain functional networks.

### Developmental refinement of opposing co-fluctuation patterns along the SA axis

The adult findings revealed amplitude-dependent configurations of co-fluctuation scores along the SA axis. To investigate how these patterns develop during childhood and adolescence, we analyzed the HCP development dataset (6-18 years old) by averaging co-fluctuation score maps across participants within the same age groups. The two clusters identified in adults were detectable as early as age 6 and persisted across development (Fig. 4A). Importantly, the distinction between the two clusters became more pronounced with age, as indicated by increasingly divergent distributions of the correlations between co-fluctuation score maps of different amplitudes (Fig. 4B). Furthermore, the amplitude-dependent alignment of co-fluctuation score maps with the SA axis exhibited progressive convergence toward adult profiles, particularly at the trough (high amplitude) and peak (intermediate amplitude), reaching adult-equivalent values by late adolescence (r_age18, adult_ = 0.98, r_age6, adult_ = 0.72, Spearman’s rho; Fig. 4, C and D). These results suggest that spatial patterns of regional co-fluctuation become more defined and specialized throughout development.

**Fig. 4.**
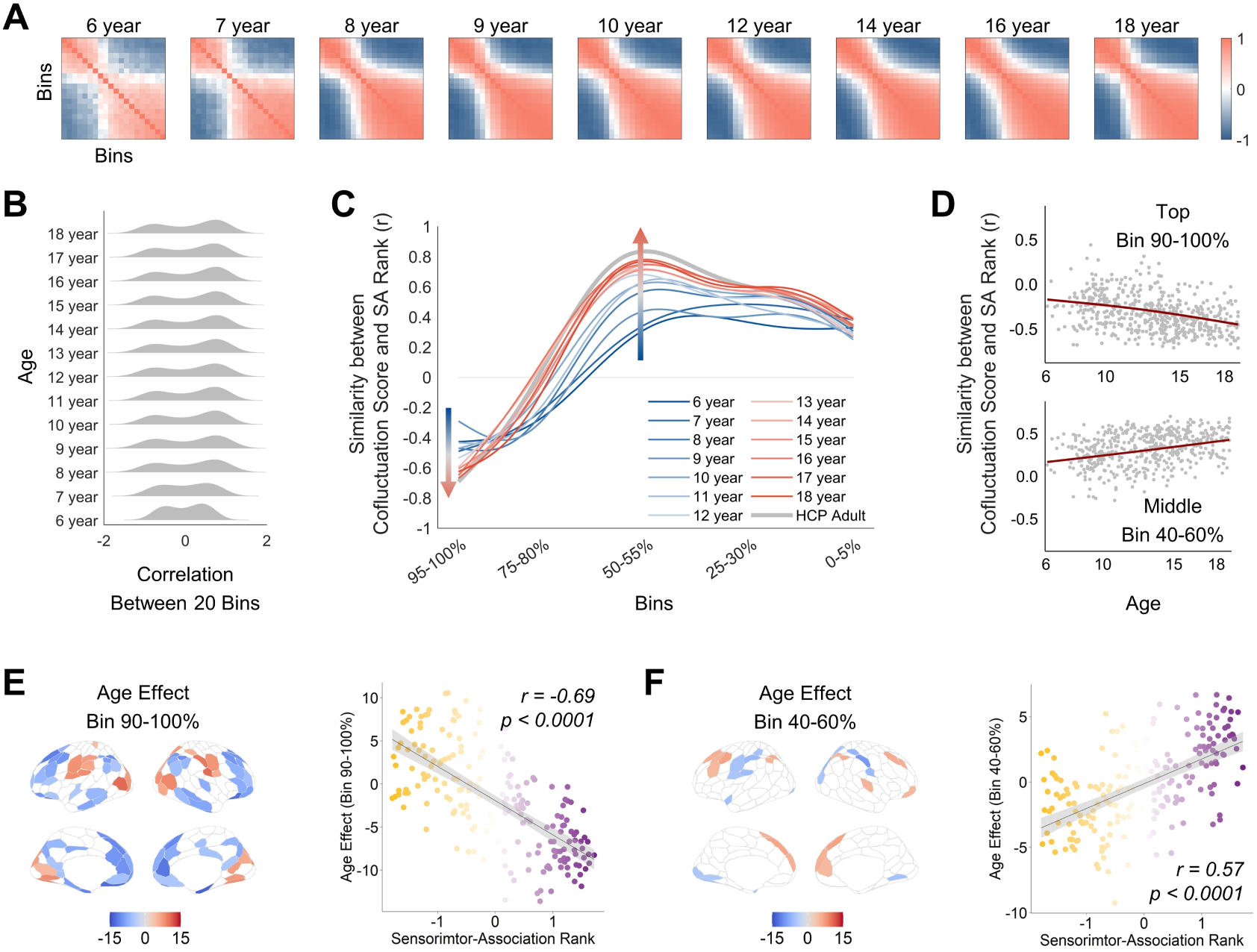
Developmental refinement of amplitude-dependent co-fluctuation hierarchy. (**A**) Correlations of co-fluctuation score maps across 20 amplitude bins from 6 to 18 years old. (**B**) Distributions of correlations of co-fluctuation score maps across 20 bins, revealing increasing dissociation of two clusters from 6 to 18 years old. (**C**) Trajectories of amplitude-dependent alignment between co-fluctuation score maps and SA rank map approach adult-like pattern (grey line) from 6 to 18 years old. To facilitate visualization, lines were smoothed and the original figure is provided in fig. S4. (**D**) The similarities between SA ranks and co-fluctuation score maps of individuals at high and intermediate amplitude bins. Each point represents an individual participant. (**E**-**F**) Left: Age-related effects on co-fluctuation scores at high- and intermediate-amplitude bins (p < 0.005, BHFDR corrected). Right: SA ranks predict age-related effects on co-fluctuation scores at high and intermediate amplitude bins.

To identify key brain regions associated with the development of regional co-fluctuation patterns, we examined age-related effects on regional co-fluctuation scores at high (90-100%) and intermediate (40-60%) amplitudes separately. At high amplitudes, regions within the sensorimotor and visual networks exhibited significant age-related increases in co-fluctuation scores, whereas regions within the default mode and control networks showed significant age-related decreases (Fig. 4E; table S1). Conversely, these regions displayed the opposite developmental effects at intermediate amplitudes (Fig. 4F; table S2). These findings suggest that the development of spatial differentiation of co-fluctuation scores unfolds along the SA axis in opposite directions, ultimately shaping the hierarchical organization of functional networks. It highlights the developmental mechanisms underlying the functional organization of the cerebral cortex.

### State-dependent modulation of regional co-fluctuation patterns

Previous section reveals the long-term changes of two opposite spatial patterns of regional co-fluctuation. Whether these patterns can change in short-term remains unclear. To address this, we investigated state modulation by comparing resting-state and movie-watching conditions in the HCP 7T fMRI dataset. The trajectories of co-fluctuation scores across the global amplitudes in both conditions were replicated in 7T dataset compared to previous results from 3T dataset (Fig. 5A).

**Fig. 5.**
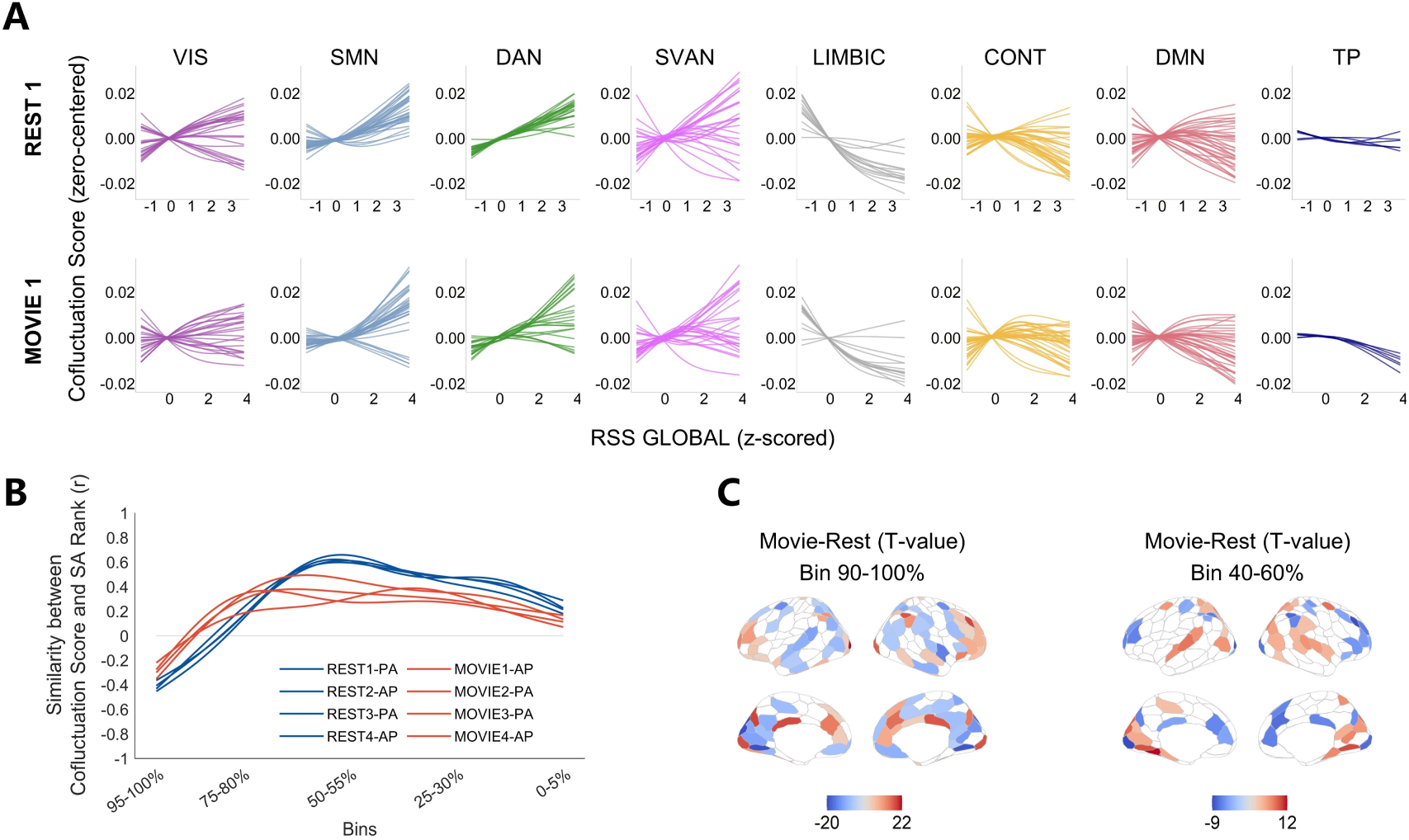
Naturalistic stimuli modulate spatial patterns of co-fluctuation scores. (**A**) Amplitude-dependent trajectories of co-fluctuation scores of seven canonical functional networks under resting-state and movie-watching conditions, derived from HCP 7T fMRI dataset. (**B**) Spatial alignments of SA ranks and co-fluctuation scores across amplitude bins in four test-retest scans during resting-state and movie-watching conditions. (**C**) Significant differences in regional co-fluctuation scores between resting-state and movie-watching conditions at high-amplitude bins (left) and intermediate-amplitude bins (right) (p < 0.05, BHFDR corrected).

However, naturalistic stimulation attenuated the alignment of co-fluctuation scores with the SA axis particularly at the high and middle amplitudes (Fig. 5B), suggesting stimulus-driven hierarchy rebalancing. Regional specificity analysis unveiled the underlying basis of this misalignment (Fig. 5C; tables S3 and S4). Sensory-processing regions (parieto-occipital/superior temporal cortices) showed significantly reduced co-fluctuation scores at high-amplitudes and enhanced that at mid-amplitudes in movie-state compared to resting-state. Conversely, high-order functioning regions (prefrontal/cingulate cortices) displayed mirror-symmetric patterns, evidencing co-fluctuation competition. Notably, the primary visual cortex exhibited a distinct state-dependent pattern. At high amplitudes, its co-fluctuation score was significantly higher in the movie-watching state, likely reflecting visual stimuli-driven processing. At middle amplitudes, its co-fluctuation score was significantly higher during the resting state, indicating suppression of the co-fluctuation of the primary visual cortex during movie-watching.

Collectively, these findings demonstrate rapid reconfiguration of co-fluctuation patterns between brain states, particularly in response to naturalistic stimuli, suggesting a dynamic modulation of functional organization that rebalance external sensory processing against internal cognitive demands.

### Sensitivity and replication analyses

We have done extended analyses to make sure our findings are reliable and replicable. First, addressing the contentious methodological issue of global signal regression, we demonstrated that the global signal is not directly related to global RSS (fig. S1). Critically, our findings are robust to the exclusion of global signals or not (fig. S3). Given the shorter scan duration in the HCP development dataset compared to the HCP dataset, we repeated the analysis with 10 bins on developmental dataset. The age-related findings remained consistent with that observed using 20 bins (fig. S4). Next, we replicated the main findings across independent datasets: 7T resting-state data recapitulated 3T-derived findings (Fig. 5A; fig. S6), while independent developmental dataset replicated age-related findings (see details in Materials and Methods, and results in fig. S5). This convergent evidence across MRI protocols, preprocessing strategies and datasets substantiates that the observed regional co-fluctuation patterns reflect intrinsic neurobiological profiles rather than methodological artifact.

## Discussion

Our study reveals an amplitude-stratified cortical dynamic architecture where spontaneous co-fluctuation patterns reorganized between functional systems along the sensorimotor-association axis (Fig. 6). Moving beyond conventional focus on high-amplitude events, our work demonstrate that dynamic co-fluctuation patterns across amplitudes are organized by conserved hierarchical principles: sensorimotor regions dominate global co-fluctuations during high-amplitude states, while associative/limbic cortices preferentially engage during mid-low amplitude regimes. This dual-dominance co-fluctuation patterns evolve during development, becoming more aligned with adult-like functional organization. Furthermore, the state-dependent modulation of these patterns under naturalistic stimulation demonstrates their adaptive reconfiguration capacity corresponding to cognitive and perceptual demands. This amplitude-stratified framework bridges transient dynamic interaction with stable cortical hierarchy, offering insights into how intrinsic functional networks are dynamically organized.

**Fig. 6.**
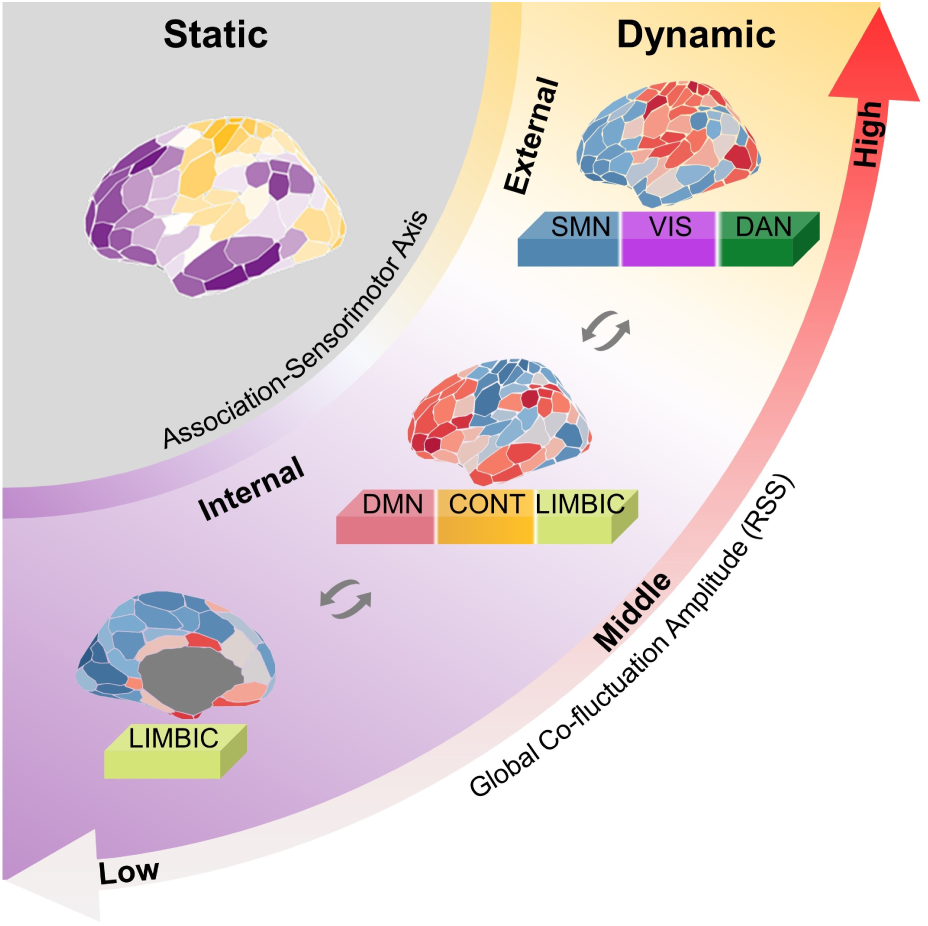
A schematic representation of the strong association between static hierarchy and dynamic co-fluctuations across high, middle and low co-fluctuation amplitudes. From static to dynamic, the hierarchy between sensorimotor and association regions remains evident, with two opposing co-fluctuation score patterns anchored at high and middle-to-low amplitudes, respectively. This underscores that dynamic co-fluctuations are intrinsically configured along the SA axis and are balanced between internal and external processing.

### Dynamic stratification of cortical interactions

Our work introduced amplitude-dependent state stratification as novel organizational principle of dynamic networks. Brain activity is ever-changing, involving dynamic interactions across functional networks that adapt us to the surrounding changing world. While conventional functional connectivity captures time-averaged coordination between regions, our amplitude-resolved decomposition demonstrates how transient co-fluctuations assemble into dynamic network states ^33^. Co-fluctuation provides complementary information beyond traditional FC metrics ^30,34,45,46^. However, prior research predominantly focused on high-amplitude co-fluctuations ^26,28,47^, neglecting low-amplitude counterparts that constitute the majority of brain activity ^9,46^. By implementing co-fluctuation score normalization, we overcome the amplitude confound to compare regional co-fluctuation patterns across different amplitude-states. Our findings revealed that regional co-fluctuations had systematically changing patterns with global amplitude. The primary and secondary amplitude effects exhibited a hierarchical distribution of three systems. Sensorimotor systems exhibited near-linear amplification of coupling with rising global amplitudes, implying their role in driving global co-fluctuation ^48^. High-order systems, especially DMN and CONT, followed inverted U-shaped trajectories, peaking at intermediate amplitudes. Limbic system demonstrated amplitude-dependent decoupling, potentially reflecting their role in dominating low-amplitude co-fluctuation. These findings delineate a tripartite amplitude-dependent reorganization of functional networks.

### Amplitude-mediated state transitions along cortical hierarchy

Our discovery of amplitude-dependent transition in regional co-fluctuation patterns unveils how the SA axis orchestrates the dynamic reconfiguration of functional interaction. This axis represents a critical organizational principle of the human cortex ^41–44^, characterized by a hierarchy from primary sensorimotor areas to high-order association areas. It conserved across the structure ^39^, function ^44^, development ^49^, and evolution ^50^ of human cerebral cortex. The functional dynamics also follow the cortical hierarchy ^22,23,51–53^. Recent dynamic modeling studies demonstrate that incorporating SA-axis topology recapitulates empirical dynamic FC ^48,54^. Hierarchical dynamics may represent an intrinsic organizational principle of brain function ^35,55,56^. It suggests our observed amplitude-dependent transitions of co-fluctuation emerge from intrinsic organization principles along the SA gradient.

While prior work identified temporal transitions between recurring functional states ^57–60^, our amplitude-resolved analysis reveals these represent meta-states of a continuum governed by SA-axis dynamics from high to middle-low amplitudes ^22^. High-amplitude states were dominated by co-fluctuation of sensorimotor systems, whereas mid-low amplitude phases engaged interaction of association/limbic systems. Esfahlani et al. (2020) found similar S-A pattern with our findings at high amplitude ^26^, but without investigating other amplitudes. The dynamic transition of co-fluctuation implies that, under the overarching constraint of the SA axis, complementary and flexible co-fluctuation patterns exist within the human cerebral cortex. High-amplitude states facilitate processing rapid-changing external information. Whereas, mid-low amplitude phases may process internal information including complex cognition and emotion.

### Developmental refinement of hierarchical dynamics

Previous studies have proposed that the development of functional connectivities across cortices unfolds along SA-axis hierarchy, with increasing segregation between sensorimotor and association regions ^41,42,49,61–66^. Here, we showed that the two opposing co-fluctuation patterns gradually became more aligned with the SA axis from childhood to adolescence – sensorimotor networks strengthening high-amplitude coupling while association cortices enhance mid-low amplitude synchronization. It exemplifies how SA-axis hierarchy scaffolds the development of brain dynamics. The developmental pattern of co-fluctuation scores at high amplitudes is consistent with existing research findings ^41,42,63,65–67^. However, the development of co-fluctuation scores at middle-to-low amplitudes is less discussed. Previous research has suggested that system segregation is crucial for cognitive ^24^, language ^68^, and task performance ^69^ processes. Our findings suggest that these opposing co-fluctuation patterns may facilitate information transmission processes between bottom-up sensory processing and top-down cognitive control, supporting increasingly complex task demands during development ^70,71^. These insights could aid in identifying atypical developmental patterns, such as those seen in neurodevelopmental disorders ^72,73^.

### Stimulus-driven co-fluctuation hierarchy reorganization

Naturalistic stimulation induced amplitude-dependent co-fluctuation patterns decoupling with SA axis. During movie-watching, sensorimotor regions exhibited suppressed co-fluctuation at high amplitudes and increased co-fluctuation at middle amplitudes, while association regions showed reverse tuning. This stimulus-driven reorganization of co-fluctuation reflects adaptive functional dynamics, where external demands transiently override intrinsic hierarchy constraints ^74–77^. Such constraint flexibility may enables rapid transitions between perceptive and complex cognitive processes to meet varying environmental challenges ^77–82^.

While our findings establish amplitude-stratified hierarchy as a core principle of dynamic functional architecture, four key frontiers warrant exploration. First, our study focused solely on conscious states. Investigating unconscious states, such as sleep ^83,84^, may further elucidate the dynamic organization of functional interactions. Second, the study examined only healthy populations. Whether psychiatric or neurodevelopmental disorders manifest atypical dynamics requires further exploration ^20,21^. Third, the neurobiological mechanisms driving amplitudes transitions were unexplored. Future research incorporating biophysical models may help uncover these mechanisms ^48,54,56,85^. Finally, how anatomical structure gives rise to the complex functions remains poorly understand ^86–88^. How structural connectome shape co-fluctuation patterns is still an open question, which would advance our understanding of structure and function coupling.

Our study unveils the human cortex employs amplitude-dependent hierarchy to organize dynamic co-fluctuations. High-amplitude states are driven by sensorimotor regions’ interaction, while mid-low amplitude regimes recruit co-fluctuation of association/limbic cortices. These opposing patterns are developmentally refined and adaptively reorganized by naturalistic stimuli, establishing the SA axis as fundamental infrastructure of brain dynamic. By resolving amplitude-dependent transitions of co-fluctuation patterns, we provide a framework for understanding how transient coordination assemble into stable functional architectures.

## Materials and Methods

### Datasets

We used three resting-state fMRI (rs-fMRI) datasets from Human Connectome Project (HCP), including 3T and 7T adults rs-fMRI datasets as well as 3T rs-fMRI developmental dataset. The Chinese Color Nest Project (CCNP) served as an independent dataset to assess the reproducibility of the developmental findings.

### Human Connectome Project (HCP) dataset

3T rs-fMRI data was from HCP S1200 release ^89^, a subset of 100 unrelated adult participants (54% females; mean age = 29.11 ± 3.67 years; age range = 22 - 36 years). The participants underwent four rs-fMRI scans with eyes open with relaxed fixation on a cross. Rs-fMRI images were acquired with a multiband gradient-echo EPI sequence (run duration = 14.33 mins, TR = 720 ms, TE = 33.1 ms, flip angle = 52°, FOV = 208 x 180 mm2, slice thickness = 2 mm, 72 slices, 2-mm isotropic voxel resolution, multiband factor = 8). All data were collected on a customized Siemens 3T Connectome Skyra scanner with a standard 32-channel head coil.

7T rs-fMRI dataset consists 182 healthy subjects (60% female; mean age = 27.53 ± 3.26 years; age range = 22 - 36 years). The participants underwent four rs-fMRI, with eyes open with relaxed fixation on a cross. Rs-fMRI images were acquired with a multiband gradient-echo EPI sequence (run duration = 16 mins, TR = 1000 ms, TE = 22.2 ms, flip angle = 45°, FOV = 208 x 208 mm2, slice thickness = 1.6 mm, 85 slices, 1.6 - mm isotropic voxel resolution, multiband factor = 5).

7T movie-watching fMRI dataset consists 182 healthy subjects (60% female; mean age = 27.53 ± 3.26 years; age range = 22 - 36 years). There are 4 movie-watching scans in total, with each 2 scans were collected after rs-fMRI scanning in session 1 and session 4. It was acquired with the same scanning protocols as rs-fMRI, except for the duration (movie 1/2/3/4 = 15.35 mins/15.3 mins/15.25 mins/15.02 mins).

The HCP study was approved by the Washington University Institutional Review Board, and all participants provided informed consent to the study protocols and procedures.

### Human Connectome Project Development (HCPD) dataset

Human connectome project development dataset ^90^ was used, which includes 652 subjects (53.9% females; mean age = 14.5 ± 4.1 years; age range = 5 - 22 years). Each participant older than 8 years underwent a total of 26-minute rs-fMRI scanning across four sessions. For the younger group (5 – 7 years), rs-fMRI scanning was divided into six sessions, with a total duration reduced to 21 mins. In this study, rs-fMRI data acquired from participants aged 6 to 18 years were analyzed, totaling 2,139 sessions from 530 individuals (53.2% females) after quality control. All participants were instructed to stay still and awake, and blink normally while looking at the fixation crosshair on the screen. The study procedures were approved by a central Institutional Review Board at Washington University in St. Louis.

### Chinese Color Nest Project (CCNP) dataset

The Chinese Color Nest Project dataset included a total of 479 participants ^91^. Each participant undergoes one, two or three visits, finally composed of a total of 864 visits. Each visit included two rs-fMRI scans. All participants were instructed to stay still and awake, and blink normally while looking at the fixation crosshair on the screen. After quality control, a total of 734 sessions were included for the subsequent analyses, comprising 403 from baseline (50.1% females; mean age = 10.66 ± 3.14 years, age range = 6 - 18.9 years), 216 from 2-th visit (52.7% females; mean age = 11.97 ± 2.76 years, age range = 7.6 - 18.9 years) and 115 from 3-th visit (55.6% females; mean age = 12.72 ± 2.51 years, age range = 8.9 - 18.9 years). For CKG sample, rs-fMRI images were acquired with EPI sequence (run duration = 7 min 45 s, TR = 2500 ms, TE = 30 ms, flip angle = 80°, 38 slices). For PEK sample, rs-fMRI images were acquired with a gradient-echo EPI sequence (run duration = 6 mins, TR = 2000 ms, TE = 30 ms, flip angle = 90°, 33 slices). CCNP was approved by the Institutional Review Board of the Institute of Psychology, Chinese Academy of Sciences. Prior to conducting the research, written informed consent was obtained from one of the participants’ legal guardians, and written assent was obtained from the participants.

### Preprocessing

All HCP and HCPD brain images were preprocessed using the ‘minimal preprocessing pipelines ^92^. The preprocessing of rs-fMRI images includes distortion and motion correction, alignment to respective T1w image with one-spline interpolation, intensity bias correction and alignment to common space using a multi-model surface registration. Then the BOLD timeseries was linearly detrended, band-pass filtered (0.008 - 0.08Hz), and 36-parameters confounds regressed including six head-motion parameters, mean signals of cerebrospinal fluid (CSF), white matters (WM), mean global signal, the derivates of above nine regressors, and the squares of above 18 parameters. A functional parcellation designed to optimize both local gradient and global similarity measures of the fMRI signals was used to define 200 regions of the cerebral cortex ^36^. For HCP dataset, timepoints with framewise displacement > 0.2 mm was excluded, and for HCPD dataset, we removed timepoints with framewise displacement > 0.3 mm.

CCNP dataset was preprocessed using Connectome Computation System (CCS) ^93^. All preprocessing scripts are available on github (https://github.com/zuoxinian/CCS). Rs- fMRI data preprocessing included the following steps: (1) dropping the first 10 s (5 TRs) for the equilibrium of the magnetic field; (2) correcting head motion; (3) slice timing; (4) despiking for the time series; (5) estimating head motion parameters; (6) aligning functional images to high resolution T1 images using boundary-based registration; (7) mitigating nuisance effects such as ICA-AROMA-derived, CSF and white matter signals; (8) removing linear and quadratic trends of the time series; (9) projecting volumetric time series to fsaverage5 cortical surface space; and (10) 6-mm spatial smoothing. Timepoints with framewise displacement > 0.3 mm were excluded and sessions with at least 60% reserved timepoints were included in this study.

### Estimating time-varying co-fluctuation

Functional connectivity (FC) is commonly calculated as the Pearson’s correlation coefficient between BOLD timeseries, which is mathematically equal to the average of temporal co-fluctuation between two regions. Therefore it can be decomposed into a series of moment-by-moment co-fluctuations, which is called edge timeseries (ETS) ^25,26,28^. E_i,j_ is defined as the element-wise product between the z-scored BOLD timeseries z_i_(t) and z_j_(t) from ROI_i_ and ROI_j_ (Fig. 1A). Each element of ETS denotes the instantaneous co-fluctuation between ROI_i_ and ROI_j_. It is worth noting that the mean of all elements in the edge timeseries is exactly equal to the corresponding FC between two ROIs. Therefore, ETS can be considered as the decomposition of FC.

### Estimating co-fluctuation score

We calculated the root sum squares (RSS) of the co-fluctuation values at each time point in ETS across different edges to quantify the co-fluctuation amplitude. The global co-fluctuation amplitude reflects the instantaneous co-fluctuation across all pairs of cortical regions at a given timepoint, while the co-fluctuation amplitude of single region captures the co-fluctuation between that region and all other brain regions.

To compare the time-varying configurations of regional co-fluctuation across different global amplitudes, we proposed a measure--co-fluctuation score, which indexes the relative co-fluctuation of a specific region. It can also be considered as the contribution of regional co-fluctuation to the global co-fluctuation. Firstly, we sorted the timepoints into 20 bins (5% timepoints per bin) based on their global co-fluctuation amplitudes, ranging from low to high. Then we calculated the mean global and regional co-fluctuation amplitudes separately within each bin. The ratio of regional to global co-fluctuation is defined as the co-fluctuation score (Eq. 1).

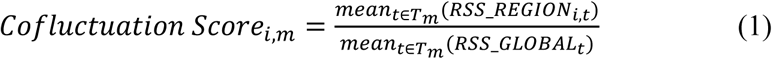

*i* represents the *i* − *t*ℎ region, m represents the *m* − *t*ℎ bin and *T_m_* means the timepoints within *m* − th bin. RSS_REGION*_i,t_* represents the RSS for the *i* − *t*ℎ region at timepoint *t* . RSS_GLOBAL*_t_* represents the RSS of whole-brain at timepoint *t* . This procedure generates a co-fluctuation score matrix (N_ROIs x N_Bins), with each element representing the contribution of a specific region to the whole-brain co-fluctuation amplitude at a given amplitude bin.

### Delineating the changes of co-fluctuation scores across global amplitudes

To explore how relative regional co-fluctuation varies across different amplitudes, we delineated the trajectory of co-fluctuation score across amplitudes for each region. We employed generalized additive models (GAMs) to flexibly model linear or non-linear relationship between regional co-fluctuation scores and global co-fluctuation amplitudes, using ‘mgcv’ package in R. GAM was fitted with co-fluctuation score as the dependent variable, the mean of global co-fluctuation amplitude within each bin as a smooth term, and sex and mean head motion as linear covariates (Eq. 2).

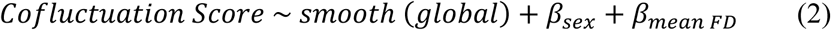

The models were fitted separately for each region using thin plate regression splines as the basis set of the smooth term and the restricted maximal likelihood approach for smoothing parameter selection. The GAM smooth term produces a spline that represents the trajectory of how regional co-fluctuation contributions to global co-fluctuation vary across global amplitudes. The significance of the association between co-fluctuation score and global amplitude was accessed by the analysis of variance that compared the full GAM model to a nested, reduced model with no smooth term. We estimated the primary amplitude effect by quantifying the difference in partial R^2^ between the full and reduced model. The direction of primary amplitude effect was determined by the sign of the averaged first derivatives of the smooth function, from which we generated a brain map of the primary amplitude effect. To further elucidate the shape of the trajectories, we calculated the averaged second-order derivatives of each trajectory, a measure we termed the second amplitude effect. Positive values indicate a U-shaped profile, while negative values correspond to an inverted U-shape pattern.

### Neurobiological underpinnings

To explore the neurobiological basis of the amplitude effect, we assessed cortical myelination and excitatory-inhibitory balance. Pearson’s correlation coefficients were calculated between the amplitude effect and two neurobiological markers: myelination and gene expression. Spin testing was used to establish statistical significance.

For cortical myelination analysis, we used the averaged T1w/T2w ratio map released by the HCP ^94^. No additional preprocessing was performed beyond averaging the myelin values within each region based on the Schaefer200x17 ^36^.

For gene expression analysis, microarray expression data and accompanying metadata were downloaded from ABHA ^95^. Group-averaged expression values for 16,088 brain specific genes across 200 (Schaefer200x17) cortical areas were used for the analyses. Finally, the difference between z-normalized gene expression values of parvalbumin (PVALB) and somatostatin (SST) was calculated.

### Meta-analysis of co-fluctuation scores

For each amplitude bin, we select the 20% brain regions with highest co-fluctuation score and then project to the 2-mm volumetric MNI152 standard space. The 20 brain volumes were input to meta-analysis (Neurosynth ^96^) and z-statistic is used to represent the association between regions and terms. Similar to previous study ^44^, 23 topic terms were select within our analysis, and the “reading” term was removed because it not captures any significant relationships at each bin.

### The relationship between co-fluctuation scores and the SA axis

The sensorimotor-association axis (SA axis) represents a cortical hierarchy that varies from unimodal (sensory) regions to transmodal (association) cortex ^41^. The cortical anatomical ^39^, functional ^44^ and evolutionary ^50^ organizations all exhibited similar patterns as SA axis.

We investigated whether the co-fluctuation scores at different amplitudes consistently align with the SA axis ^41^. Specifically, we averaged co-fluctuation score map at each amplitude bin across subject as group-level co-fluctuation score. Then we correlated the SA rank (z-scored) map with the group-level co-fluctuation score map at each amplitude bin. For the HCPD dataset, we averaged co-fluctuation score maps across subjects within each age group (ranging from 6 to 18 year) to assess whether the alignment of co-fluctuation scores with the SA axis changes progressively during development. For better visualization, the trajectories of the similarity between SA ranks and co-fluctuation scores across bins were smoothed by fitting with GAM for each age group. Raw trajectories are illustrated in fig. S4.

## Acknowledgments

**Funding:** This work is supported by the National Natural Science Foundation of China (82102134 to Y.H.). HCP data were provided by the Human Connectome Project, WU-Minn Consortium (Principal Investigators: David Van Essen and Kamil Ugurbil; 1U54MH091657) funded by the 16 NIH Institutes and Centers that support the NIH Blueprint for Neuroscience Research; and by the McDonnell Center for Systems Neuroscience at Washington University.

## Data and materials availability

The Human Connectome Project adult dataset is publicly available at https://db.humanconnectome.org. The Human Connectome Project developmental dataset is available at the National Institute of Mental Health Data Archive. Chinese Color Nest Project dataset is available at the Science Data Bank (https://doi.org/10.57760/sciencedb.07478). Code for analyzing co-fluctuation scores is available in GitHub (https://github.com/dzjin5678/co-fluctuation-scores).

## Supplementary Materials

**Fig. S1.**
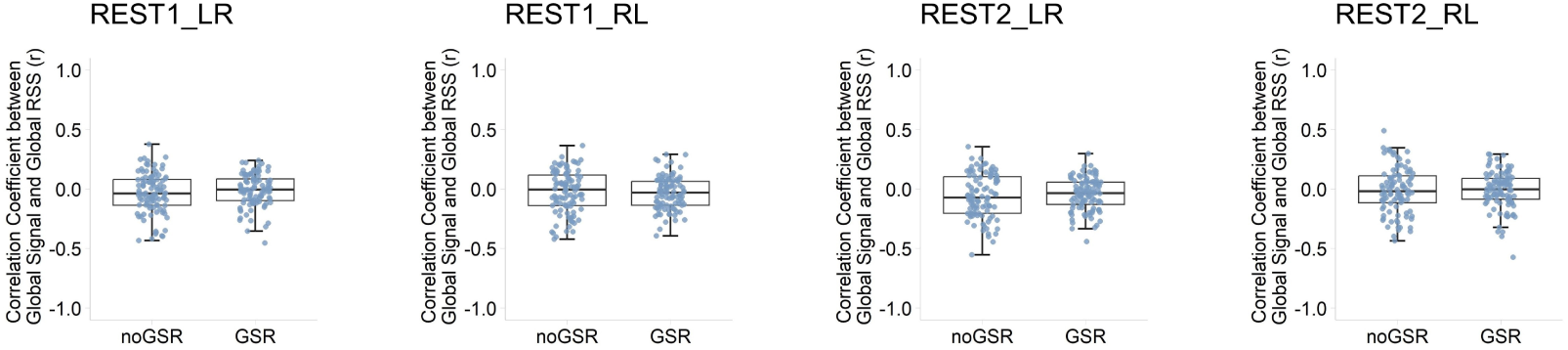
The relationship between global signal and global RSS. We calculated Pearson’s correlation coefficient between global signal and global RSS for four scan sessions of each subject. ‘noGSR’ refers to the condition that global RSS was calculated without global signal regression, whereas ‘GSR’ indicates that global RSS was calculated after global signal regression. There was no significant difference between the conditions of noGSR and GSR, which suggests global signal is not directly related to global RSS.

**Fig. S2.**
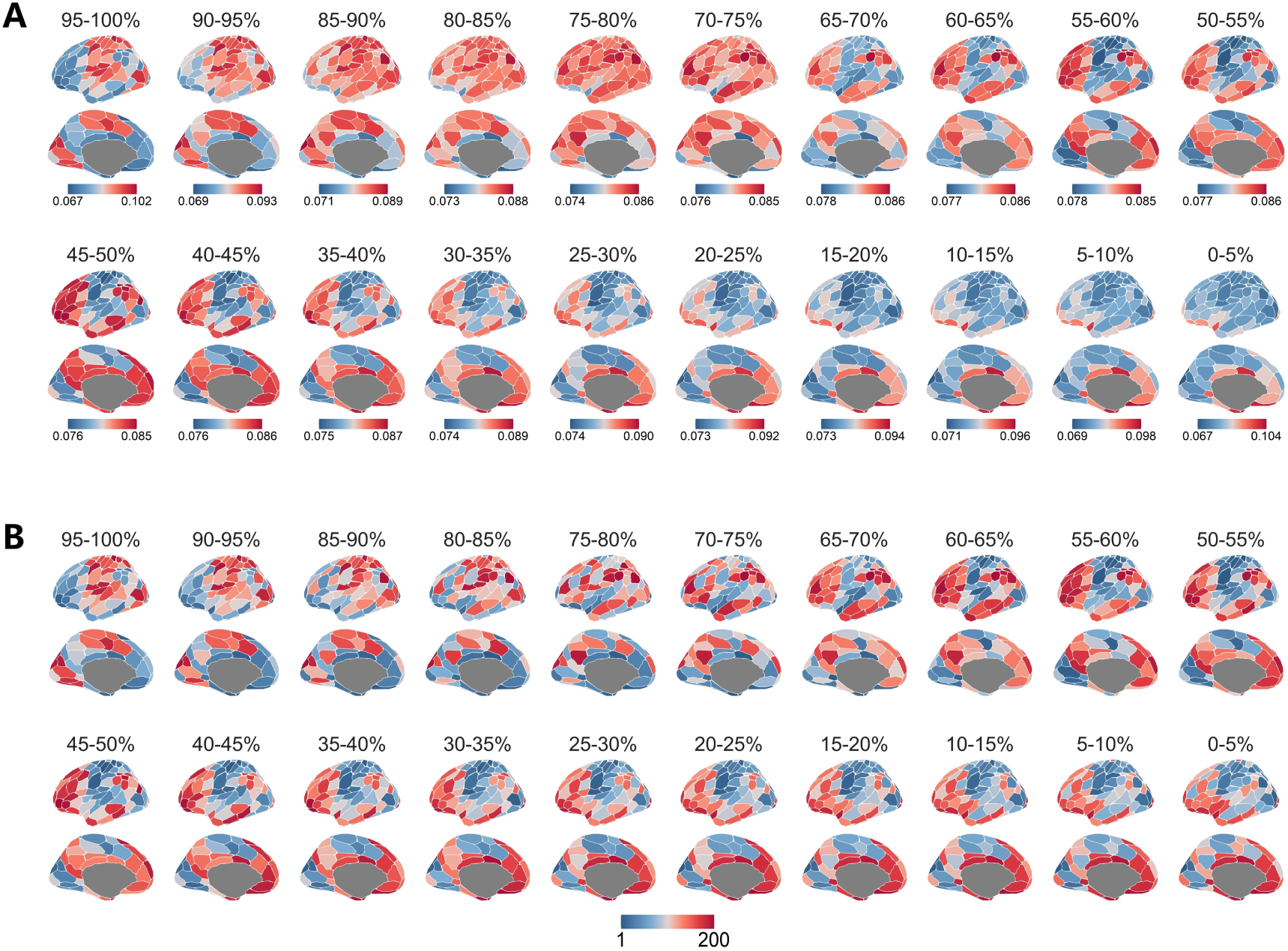
Co-fluctuation score maps for 20 amplitude bins derived from the HCP 3T dataset following global signal regression. (**A**) Raw co-fluctuation scores for each bin. (**B**) To facilitate direct comparison of the hierarchy of co-fluctuation scores in the cerebral cortex, we plotted their ranks (from 1 to 200) for all bins.

**Fig. S3.**
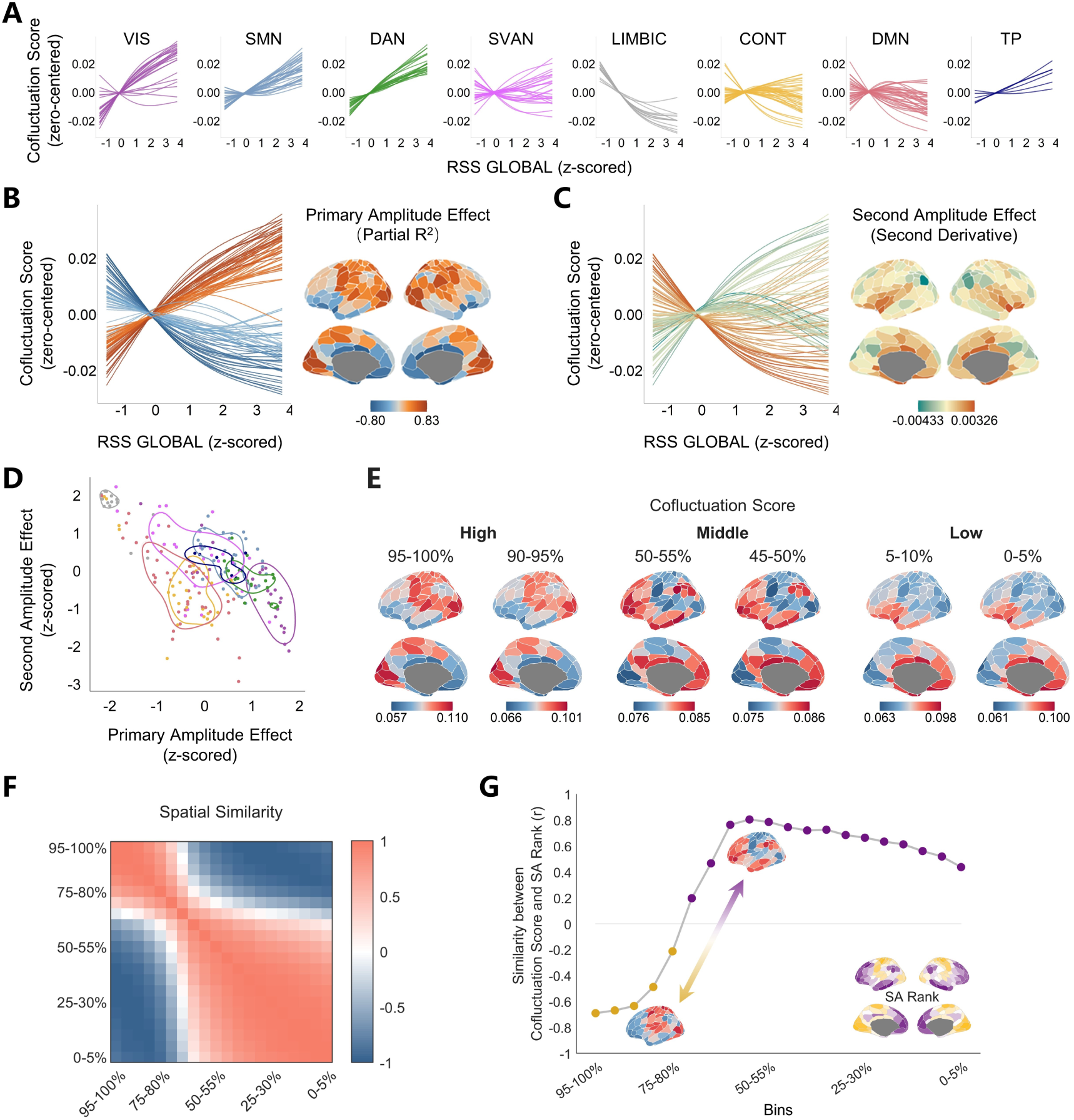
Sensitivity analysis without global signal regression. The same analysis has been conducted using the same data but preprocessed without global signal regression. The findings could be replicated as in Fig. 2. (**A**) Co-fluctuation score trajectories were grouped for eight canonical functional networks. (**B**) The primary amplitude effect (partial R^2^) summaries the overall changing trend of co-fluctuation scores with global amplitudes. (**C**) The second amplitude effect (averaged second-order derivatives) further characterizes the shape of trajectories. (**D**) The scatter plot of the primary and second amplitude effects. (**E**) The regional co-fluctuation score maps at the high, middle and low amplitude bins. (**F**) The similarity of spatial patterns of co-fluctuation scores across 20 bins (Pearson’s correlation coefficient). (**G**) The correlation between co-fluctuation score maps and SA rank map at each bin (Spearman correlation coefficient).

**Fig. S4.**
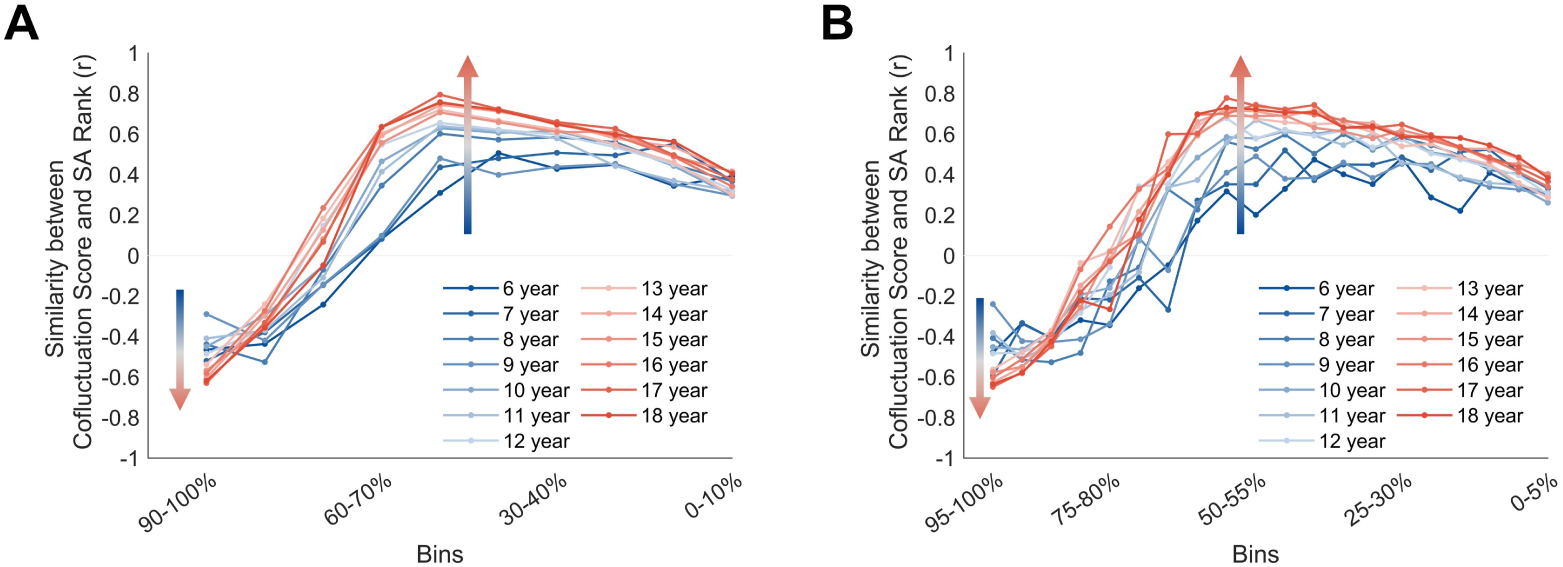
Sensitivity analysis of the maturation of similarity between SA rank and co-fluctuation score. (**A**) We reorganize all frames into 10 bins (10% timepoints per bin) according their global RSSs. In this case, the number of timepoints is nearly twice that of 20 bins. (**B**) The raw similarity trajectories corresponding to Fig. 4C are provided.

**Fig. S5.**
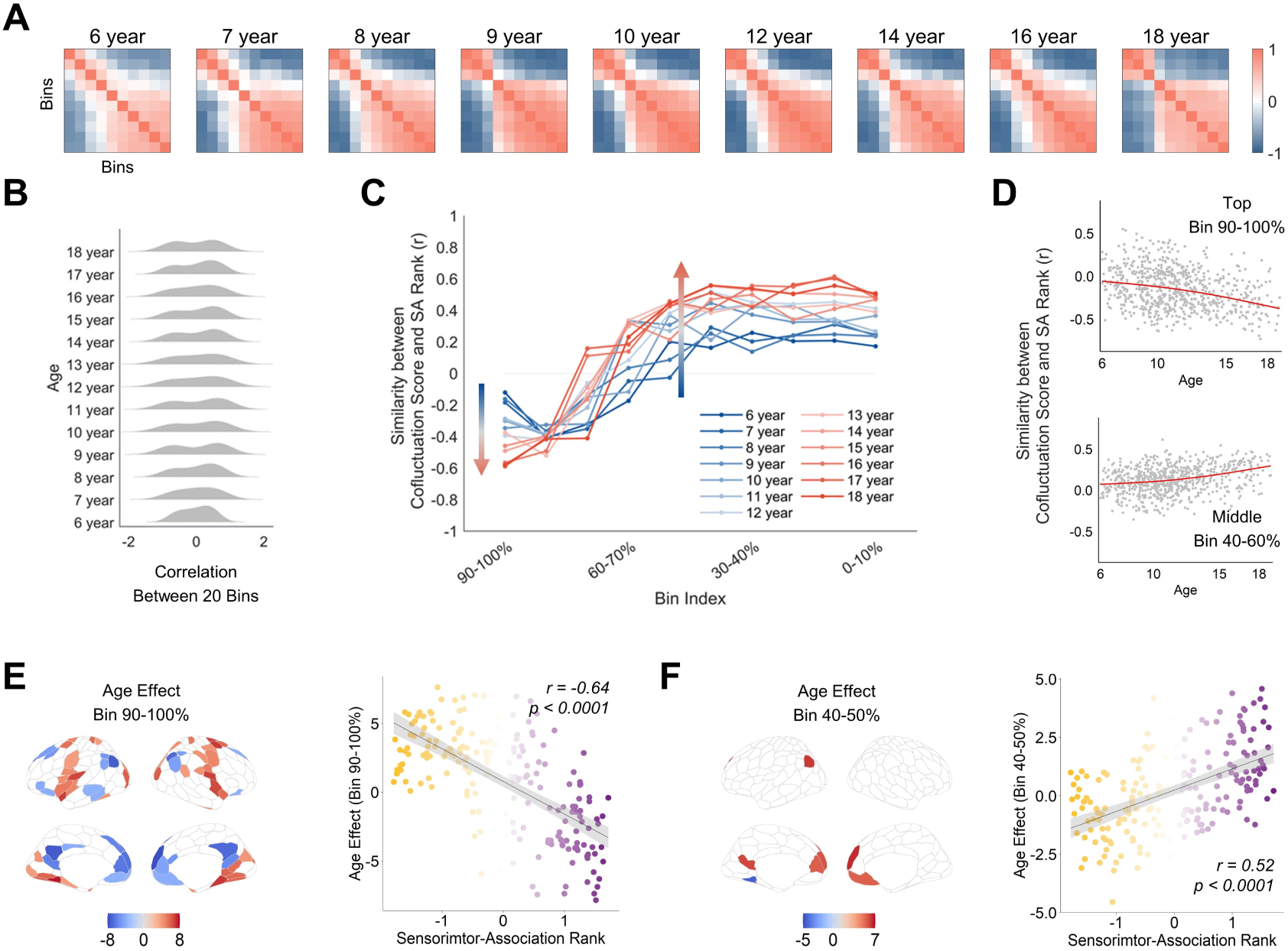
The reproducibility of developmental findings using independent dataset (CCNP). Due to the short rs-fMRI scanning duration for the CCNP dataset, we set the number of bins to10 to ensure the stability of the results. The findings could be replicated as shown in Fig. 4. (**A**) Correlations of co-fluctuation score maps across 20 amplitude bins from 6 to 18 years old. (**B**) Distributions of correlations of co-fluctuation score maps across 20 bins, revealing increasing dissociation of two clusters from 6 to 18 years old. (**C**) Trajectories of amplitude-dependent alignment between co-fluctuation score maps and SA rank map approach adult-like pattern (grey line) from 6 to 18 years old. (**D**) The similarities between SA ranks and co-fluctuation score maps of individuals at high and intermediate amplitude bins. Each point represents an individual participant. (**E**-**F**) Left: Age-related effects on co-fluctuation scores at high- and intermediate-amplitude bins (p < 0.005, BHFDR corrected). Right: SA ranks predict age-related effects on co-fluctuation scores at high and intermediate amplitude bins.

**Fig. S6.**
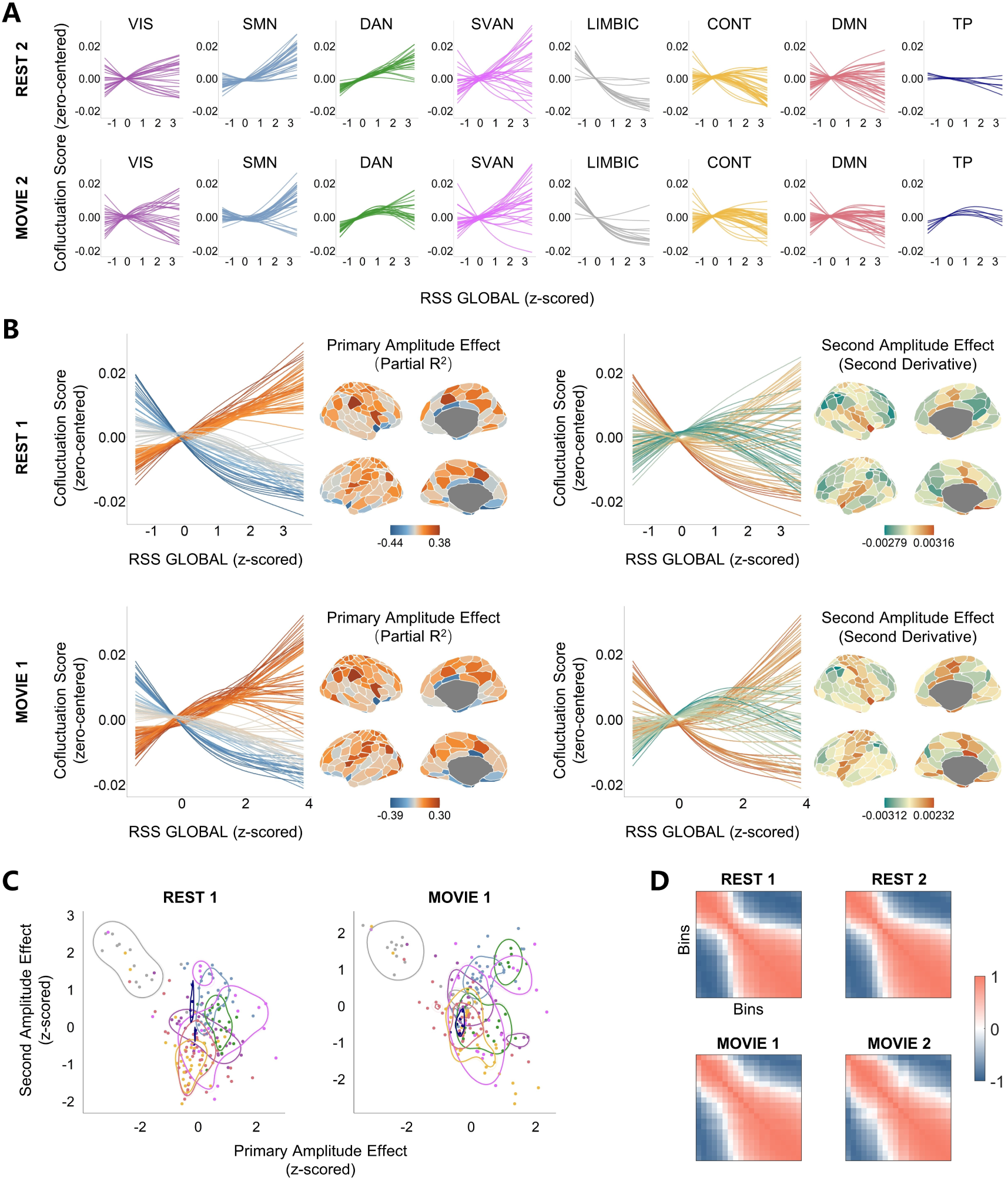
Replication of the findings on adults using 7T fMRI data. (**A**) Regional trajectories of co-fluctuation scores of eight canonical functional networks during the second resting-state and movie-watching scans, derived from a 7T fMRI dataset. (**B**-**C**) The primary and second amplitude effects during the first resting-state and movie-watching sessions. (**D**) Similarities of co-fluctuation score maps across 20 amplitude bins under each of two resting-state and move-watching sessions.

**Table S1.** Age effects of regional co-fluctuation score at high-amplitude bin (90-100%).

| Positive Age Effect |  |  | Negative Age Effect |  |  |
| --- | --- | --- | --- | --- | --- |
| ROIName | T | p | ROIName | T | p |
| RH_SomMotB_S2_4 | 10.6 | 0 | LH_LimbicB_OFC_1 | -13.6 | 0 |
| LH_DorsAttnA_ParOcc_1 | 10.5 | 0 | RH_LimbicA_TempPole_3 | -13.5 | 0 |
| RH_DorsAttnB_PostC_1 | 9.6 | 0 | LH_LimbicB_OFC_2 | -12.7 | 0 |
| RH_DorsAttnA_ParOcc_1 | 8.7 | 0 | RH_LimbicB_OFC_2 | -12.5 | 0 |
| LH_VisCent_ExStr_1 | 8.7 | 0 | RH_LimbicB_OFC_1 | -12.4 | 0 |
| LH_VisCent_ExStr_5 | 8.6 | 0 | RH_DefaultA_PFCm_1 | -11.9 | 0 |
| LH_SomMotB_S2_3 | 8.5 | 0 | LH_DefaultA_PFCd_1 | -11.3 | 0 |
| LH_VisPeri_ExStrInf_2 | 8.4 | 0 | LH_LimbicA_TempPole_1 | -11.2 | 0 |
| LH_SomMotB_S2_2 | 8.2 | 4.44E-16 | RH_ContB_PFCmp_1 | -10.6 | 0 |
| RH_VisCent_ExStr_1 | 8.1 | 8.88E-16 | LH_DefaultB_PFCd_3 | -10.5 | 0 |
| LH_SomMotB_Cent_1 | 8.0 | 1.78E-15 | LH_ContA_IPS_1 | -10.3 | 0 |
| RH_VisCent_ExStr_4 | 7.9 | 5.77E-15 | RH_LimbicA_TempPole_2 | -10.2 | 0 |
| RH_SomMotB_Cent_1 | 7.8 | 6.22E-15 | RH_ContB_IPL_2 | -10.0 | 0 |
| RH_VisCent_ExStr_5 | 7.8 | 7.99E-15 | LH_DefaultA_PFCm_3 | -9.9 | 0 |
| RH_DorsAttnA_SPL_4 | 7.4 | 1.38E-13 | LH_DefaultB_PFCd_4 | -9.8 | 0 |
| RH_VisPeri_ExStrInf_1 | 7.4 | 2.23E-13 | RH_ContB_PFCld_2 | -9.7 | 0 |
| LH_VisCent_ExStr_4 | 7.4 | 2.33E-13 | LH_DefaultB_PFCv_2 | -9.7 | 0 |
| LH_VisPeri_ExStrSup_2 | 7.1 | 1.28E-12 | LH_ContA_PFCI_3 | -9.7 | 0 |
| LH_DorsAttnB_PostC_1 | 7.1 | 1.81E-12 | LH_SalVentAttnB_PFCmp_1 | -9.5 | 0 |
| RH_VisPeri_ExStrSup_1 | 7.1 | 1.84E-12 | RH_ContA_IPS_1 | -9.5 | 0 |

**Table S2.** Age effects of regional co-fluctuation score at middle-amplitude bin (40- 60%).

| Positive Age Effect |  |  | Negative Age Effect |  |  |
| --- | --- | --- | --- | --- | --- |
| ROIName | T | p | ROIName | T | p |
| RH_ContB_PFCmp_1 | 6.7 | 2.86E-11 | LH_VisCent_ExStr_1 | -5.2 | 2.72E-07 |
| LH_ContA_PFCI_3 | 6.7 | 3.15E-11 | LH_VisCent_ExStr_2 | -5.0 | 7.84E-07 |
| LH_DefaultB_PFCd_3 | 6.4 | 1.40E-10 | LH_VisCent_Striate_1 | -4.7 | 2.19E-06 |
| LH_ContA_IPS_1 | 5.9 | 3.08E-09 | LH_VisCent_ExStr_3 | -4.4 | 1.12E-05 |
| RH_ContB_PFCId_2 | 5.9 | 3.71E-09 | LH_VisCent_ExStr_4 | -4.4 | 1.37E-05 |
| RH_ContB_IPL_2 | 5.7 | 1.21E-08 | LH_VisCent_ExStr_5 | -4.5 | 7.74E-06 |
| RH_ContB_PFCId_1 | 5.6 | 1.95E-08 | LH_VisPeri_ExStrInf_1 | -3.2 | 0.0016 |
| LH_DefaultB_PFCd_1 | 5.5 | 4.32E-08 | LH_VisPeri_ExStrInf_2 | -4.6 | 4.90E-06 |
| LH_DefaultA_PFCd_1 | 5.5 | 4.47E-08 | LH_SomMotA_6 | -3.1 | 0.0018 |
| RH_DefaultA_PFCm_3 | 5.4 | 7.37E-08 | LH_SomMotB_S2_2 | -5.0 | 5.01E-07 |
| LH_ContB_PFCI_1 | 5.4 | 9.45E-08 | LH_SomMotB_S2_3 | -5.8 | 7.46E-09 |
| LH_DefaultB_PFCd_4 | 5.3 | 1.44E-07 | LH_SomMotB_Cent_1 | -5.2 | 2.41E-07 |
| RH_SomMotB_Aud_1 | 5.2 | 1.90E-07 | LH_DorsAttnA_TempOcc_2 | -4.0 | 5.44E-05 |
| LH_ContB_IPL_1 | 5.1 | 3.52E-07 | LH_DorsAttnA_ParOcc_1 | -3.7 | 0.0002 |
| RH_ContB_PFCIv_1 | 4.9 | 8.27E-07 | LH_DorsAttnA_SPL_3 | -3.2 | 0.0013 |
| RH_ContA_IPS_1 | 4.8 | 1.32E-06 | LH_DorsAttnB_PostC_1 | -6.4 | 2.32E-10 |
| LH_ContA_PFCI_1 | 4.8 | 1.55E-06 | LH_DorsAttnB_PostC_2 | -5.3 | 1.19E-07 |
| RH_DefaultA_PFCm_1 | 4.8 | 1.79E-06 | LH_DorsAttnB_PostC_3 | -3.0 | 0.0024 |
| RH_SalVentAttnB_PFCIv_1 | 4.5 | 6.50E-06 | LH_LimbicA_TempPole_3 | -5.4 | 5.63E-08 |
| LH_ContA_IPS_2 | 4.5 | 6.98E-06 | LH_LimbicA_TempPole_4 | -5.6 | 2.72E-08 |

**Table S3.** Differences of regional cofluctuation score between movie and rest conditions at high-amplitude bin (90-100%).

| Movie > Rest |  |  | Movie < Rest |  |  |
| --- | --- | --- | --- | --- | --- |
| ROIName | T | p | ROIName | T | p |
| LH_VisCent_ExStr_3 | 22.1 | 4.49E-51 | LH_VisPeri_ExStrSup_2 | -19.6 | 1.88E-44 |
| LH_ContC_pCun_1 | 18.8 | 3.43E-42 | LH_VisPeri_ExStrInf_2 | -19.5 | 6.23E-44 |
| RH_ContB_PFCld_1 | 18.6 | 1.47E-41 | RH_VisPeri_ExStrInf_1 | -19.0 | 1.34E-42 |
| LH_ContC_Cingp_1 | 18.3 | 1.08E-40 | RH_VisPeri_ExStrSup_3 | -16.6 | 6.43E-36 |
| LH_VisCent_Striate_1 | 18.0 | 6.33E-40 | RH_SalVentAttnA_Ins_2 | -14.7 | 2.65E-30 |
| RH_VisCent_Striate_1 | 17.7 | 4.21E-39 | LH_DorsAttnB_FEF_1 | -13.6 | 3.90E-27 |
| RH_ContB_IPL_2 | 17.5 | 1.96E-38 | RH_DorsAttnB_FEF_1 | -13.0 | 1.56E-25 |
| RH_ContC_Cingp_1 | 17.3 | 6.75E-38 | RH_VisPeri_ExStrSup_2 | -12.7 | 1.96E-24 |
| RH_VisCent_ExStr_3 | 16.6 | 6.05E-36 | RH_VisPeri_ExStrSup_1 | -12.1 | 7.64E-23 |
| RH_SalVentAttnB_PFCmp_1 | 15.3 | 2.89E-32 | LH_SomMotB_Aud_3 | -10.9 | 2.03E-19 |
| RH_ContC_pCun_1 | 14.8 | 1.26E-30 | LH_DorsAttnA_SPL_2 | -10.9 | 2.11E-19 |
| LH_SalVentAttnB_PFCmp_1 | 14.3 | 3.37E-29 | LH_VisPeri_StriCal_1 | -10.8 | 5.06E-19 |
| RH_SalVentAttnB_Ins_1 | 13.7 | 1.93E-27 | RH_DorsAttnB_PostC_3 | -10.5 | 4.90E-18 |
| LH_ContB_PFClv_2 | 13.3 | 2.61E-26 | LH_VisPeri_ExStrSup_1 | -10.3 | 1.43E-17 |
| LH_ContA_IPS_1 | 12.1 | 7.73E-23 | RH_DorsAttnA_SPL_3 | -9.5 | 2.85E-15 |
| RH_SalVentAttnB_IPL_1 | 12.1 | 8.27E-23 | LH_SomMotB_Aud_2 | -9.2 | 1.38E-14 |
| RH_LimbicA_TempPole_4 | 12.1 | 1.02E-22 | LH_DefaultA_pCunPCC_1 | -9.2 | 1.91E-14 |
| LH_VisCent_ExStr_1 | 11.4 | 7.52E-21 | RH_LimbicB_OFC_3 | -9.2 | 2.36E-14 |
| LH_ContA_PFClv_1 | 11.0 | 1.15E-19 | RH_ContB_Temp_1 | -9.1 | 4.31E-14 |
| LH_LimbicA_TempPole_2 | 9.8 | 3.38E-16 | LH_DefaultA_IPL_1 | -8.9 | 8.80E-14 |

**Table S4.** Differences of regional cofluctuation score between movie and rest conditions at middle-amplitude bin (40-60%).

| Movie > Rest |  |  | Movie < Rest |  |  |
| --- | --- | --- | --- | --- | --- |
| ROIName | T | p | ROIName | T | p |
| LH_VisPeri_ExStrInf_1 | 11.9 | 2.87E-22 | LH_VisCent_Striate_1 | -9.0 | 5.09E-14 |
| RH_VisPeri_ExStrInf_1 | 10.1 | 4.61E-17 | RH_ContB_PFCld_1 | -8.8 | 1.91E-13 |
| LH_VisPeri_ExStrInf_2 | 9.9 | 1.97E-16 | LH_VisCent_ExStr_3 | -8.6 | 9.83E-13 |
| LH_VisPeri_ExStrSup_2 | 9.2 | 2.27E-14 | RH_ContB_IPL_2 | -7.6 | 2.91E-10 |
| RH_DorsAttnB_FEF_1 | 8.7 | 5.39E-13 | RH_VisCent_Striate_1 | -7.3 | 1.42E-09 |
| RH_VisPeri_ExStrSup_3 | 8.6 | 8.83E-13 | RH_SalVentAttnB_PFCmp_1 | -7.3 | 2.00E-09 |
| LH_SomMotB_Aud_2 | 8.4 | 1.99E-12 | RH_VisCent_ExStr_3 | -7.0 | 1.25E-08 |
| LH_DorsAttnA_SPL_2 | 8.3 | 3.62E-12 | LH_ContA_IPS_1 | -6.5 | 1.61E-07 |
| RH_DorsAttnA_SPL_1 | 7.9 | 4.08E-11 | RH_SalVentAttnB_PFCI_1 | -6.5 | 1.90E-07 |
| RH_VisPeri_ExStrSup_1 | 7.6 | 2.43E-10 | RH_DefaultA_PFCm_1 | -6.3 | 3.84E-07 |
| RH_DefaultC_PHC_1 | 7.6 | 4.02E-10 | LH_ContA_PFClv_1 | -6.2 | 6.03E-07 |
| LH_SomMotB_Aud_1 | 7.5 | 6.71E-10 | LH_ContB_PFClv_2 | -6.2 | 7.43E-07 |
| RH_DorsAttnB_PostC_3 | 7.2 | 2.82E-09 | RH_ContB_PFCmp_1 | -6.2 | 9.43E-07 |
| LH_SomMotB_Aud_3 | 7.1 | 5.55E-09 | RH_SalVentAttnB_IPL_1 | -6.1 | 1.15E-06 |
| RH_SomMotB_Aud_2 | 7.1 | 6.19E-09 | RH_ContC_Cingp_1 | -6.1 | 1.25E-06 |
| LH_DorsAttnA_ParOcc_1 | 7.0 | 9.57E-09 | LH_ContC_pCun_1 | -6.1 | 1.27E-06 |
| RH_VisPeri_ExStrSup_2 | 6.8 | 2.37E-08 | LH_SalVentAttnB_PFCmp_1 | -6.0 | 1.99E-06 |
| LH_DorsAttnB_FEF_1 | 6.8 | 2.52E-08 | RH_ContC_pCun_1 | -5.6 | 1.50E-05 |
| RH_DorsAttnA_SPL_4 | 6.7 | 5.73E-08 | RH_DefaultB_PFCv_1 | -5.2 | 9.37E-05 |
| RH_SomMotB_Aud_1 | 6.5 | 1.25E-07 | LH_SalVentAttnB_PFCI_1 | -5.2 | 0.0001 |

